# Phase-noise tolerant fully-coherent optical coherence tomography angiography for high-contrast 3D microvasculature mapping

**DOI:** 10.64898/2026.09.30.755749

**Authors:** Tianhui Jie, Sebastián Ruiz Lopera, Brett E. Bouma, Néstor Uribe-Patarroyo

## Abstract

Optical coherence tomography angiography (OCTA) has been extensively explored for the detection of pathological retinal changes arising in various ophthalmological diseases. Due to phase instability of conventional systems, most OCTA signal processing methods to date have relied on intensity-based averaging, despite the resulting lower contrast and dynamic range compared to coherent averaging approaches. In this work, we (1) present a novel computation pipeline that enables fully-coherent processing in any phase-stable or phase-unstable system, and (2) demonstrate the first angiography metric with integrated signal-to-noise ratio (SNR) correction and coherent spatiotemporal ensemble averaging effective in any OCT system. Our method does not rely on 2D *en face* projections or binarization to produce high-contrast *en face* views and it significantly enhances the contrast-to-noise ratio of computed OCTA volumes, particularly in regions with reduced SNR levels. Our OCTA volumes also have high cross-sectional quality, enabling the use of single-depth *en face* views and the localization of capillary paths in the axial dimension. We can now identify capillaries below the retinal pigment epithelium, near the lamina cribrosa in the optic cup, which was not previously possible due to low SNR. We also conducted an analysis of strategies for bias reduction in OCTA and developed the optimal method utilizing coherent spatiotemporal ensemble averaging. Because our method leverages coherent averaging across all dimensions, it provides flexibility in shaping the averaging kernel to target specific workflows. The value of this additional degree of freedom is illustrated in our vessel tracing investigations, in which the depth resolution was maximized by limiting coherent ensemble averaging to the two lateral dimensions even in presence of strong phase noise, enabling the identification of 3D features such as overlapping capillary paths.

## 1. Introduction

Optical coherence tomography angiography (OCTA) is a non-invasive imaging technique aimed at enhancing the delineation of the retinal vascular network [1, 2]. It generates depth-resolved vessel maps without requiring exogenous contrast agents, giving a significant advantage compared to dye-based fluorescein angiography when imaging to detect vascular changes arising from disease. In neovascular age-related macular degeneration, neovascularization is identified by the appearance of new/extended branches in the vessel map [3]; in proliferative diabetic retinopathy, the development of vascular occlusions manifests as disappearing patches in the vessel map [4–6]. These vascular changes can be observed at the microvascular level. Therefore, early retinal disease detection may be improved with increased capillary contrast and visibility.

In conventional OCTA, absolute phase information is unavoidably obscured by phase noise from system instabilities and subject motion. Therefore, many commercial OCTA systems [7] rely on intensity-based angiography metrics and incoherent ensemble averaging, such as split-spectrum amplitude decorrelation angiography (SSADA) [8], to avoid the impact of phase instability in the generation of OCTA volumes. Efforts have been made to include phase information in the angiography metric to increase the efficiency and realize greater angiographic contrast without requiring longer acquisition time. For example, split-spectrum amplitude phase gradient angiography (SSAPGA) [9] compounds a separately calculated phase term, and the complex differential variance (CDV) metric [10] applies coherent averaging along the axial dimension, where phase stability is intrinsic to Fourier-domain OCT.

However, practical, fully-coherent ensemble processing in presence of strong phase noise has not yet been demonstrated, presumably due to the inability to computationally remove phase noise along all dimensions. Indeed, Makita *et al*. modified their OCTA metric specifically for systems with phase jitter by limiting ensemble averaging to those dimensions acquired simultaneously and thus not affected by phase noise (i.e., the axial direction in raster-scanned Fourier-domain OCT systems) [11]. As it has been shown previously, although achieving 2D phase stability (along depth and one additional dimension) from phase unstable data is straightforward, attaining 3D or 4D phase stability is extremely challenging [12]. The use of a complex adaptive spatiotemporal kernel has also been attempted to selectively use values minimally impacted by motion-induced phase noise, but this method requires significant temporal sampling to support the model analysis [13], which is not compatible with clinically feasible scan times of a few seconds [6, 7, 14]. As a consequence, these algorithms forgo complex-valued signal fluctuations representative of flow, have significant statistical bias in correlation estimation which limits dynamic range, and restrict the effectiveness of averaging. Depending on the number of B-scan repetitions, the limitations that come with using incoherent averaging as opposed to coherent averaging are increasingly magnified [15]. We quantify these drawbacks in Section 5.1 with methods in Supplement 1, Section 1.

Prior work on implementing corrections to compensate for the contribution of noise to the decorrelation to uncover angiographic signals in regions with low signal-to-noise ratio (SNR) have not seen significant adoption. One potential reason is that the intrinsic variance in the estimation of the decorrelation requires a sufficiently large ensemble average to make the SNR correction feasible [11]. Effectively, effective SNR correction requires large ensemble sizes that are more effectively achieved with 3D/4D ensemble averaging, negating their use in systems with significant phase noise. Prior efforts have also been affected by the use of inaccurate models of the OCT noisy signal: the Rice distribution is a good fit for noisy OCT signals that are stable in time, but signals that decorrelate require a more general model [16]. In the work by Huang. *et al*., a more general model was used to mask noisy regions but it did not enable removing the contributions from noise and recalculating the actual, noiseless OCT signal decorrelation [17]. Therefore, there is a need for an SNR-corrected OCTA metric that is accurate in terms of the model used and practical implementation of large ensemble averages in any OCT system regardless of level of phase noise.

In addition to relying primarily on intensity-based OCTA metrics, standard OCTA processing protocols generally neglect the range of measured angiographic values with the use of binarization algorithms combined with the use of 2D *en face* projections to perform diagnostic analysis such as vessel area density, branchpoint density, and vessel tortuosity [14, 18, 19]. However, the continuous-valued angiographic metric, generally referred to as flux, has been show to correlate with flow speed in sufficiently small capillaries and to hold potential in diagnostics for identifying different disease stages [20, 21]. Additionally, studies have also shown the potential increase in disease specificity of metrics that take advantage of the 3D geometry of the retinal vascular network [22–26]. Therefore, improved OCTA processing should output volumes with high cross-sectional quality while maintaining its range of angiographic values.

Here, we introduce fully-coherent OCTA (FC-OCTA), a framework for computing fully-coherent OCTA in phase stable or phase unstable systems. We provide an analysis of strategies for bias reduction in OCTA, quantifying the benefits of fully-coherent spatiotemporal ensemble averaging. We demonstrate the ability to produce OCTA volumes with high angiographic contrast for retinal microvasculature **without binarization or projection** with the integration of an analytical SNR correction (A1C-SNR) to our 1^st^-order autocorrelation based contrast (A1C) angiography metric. In comparison to partially-coherent and incoherent angiography techniques such as SSADA, SSAPGA, and CDV, we are able to visualize capillary paths with increased depth localization, allowing for the identification of features such as overlapping vasculature, and at greater depths, allowing for capillary identification as deep as the lamina cribrosa in the optic cup. All data is acquired through a conventional SS-OCT system with strong phase jitter and phase noise induced by subject motion using conventional B-scan repeat protocols.

## 2. Theory

### 2.1. Enabling fully-coherent processing in highly phase-unstable systems for OCTA volume calculation

In previous investigations of noise and bias in OCT, temporal ensemble averaging has been shown to be the most efficient bias reduction strategy [15]. Works in dynamic light scattering (DLS) leverage the effectiveness of temporal averaging with long M-mode scans [27]. However, for clinical feasibility, standard OCTA imaging protocol requires short acquisition times and involves few B-scan repetitions. The next best strategy for minimizing statistical bias is spatial ensemble averaging [15]. Therefore, for OCTA applications, greatest statistical bias reduction can be achieved through 4D coherent ensemble averaging. In Section 5.1, we show results that confirm this using simulated data and widely used OCTA metrics; SSADA (entirely incoherent) and CDV (coherent axially, incoherent laterally and temporally), and our fully-coherent metric, A1C (details in Supplement 1, Section 1).

However, coherent averaging is not meaningful when OCT data is corrupted by phase noise from subject motion or system instabilities (illustrative results provided in Section 5.2), thus computationally removing phase noise is critical. At first sight, it may seem that enforcing phase stability along all four spatio-temporal dimensions is required for performing 4D coherent ensemble averaging. This demand is extremely challenging. There are methods to correct for phase noise along two orthogonal axes simultaneously, however, only in systems with negligible system instabilities and stabilized excised tissue for imaging [28]. In presence of strong phase instability, including phase jitter and subject motion, numerical corrections work only along a single axis at a time [12]. However, in the subsequent paragraphs we show that the problem can be re-framed as **computationally correcting for inter-B-scan phase noise**, thus reducing the requirement to a single-axis and enabling the use of existing 1D techniques [1, 12].

The measured OCT signal at time *t*, depth *z* and transverse position (*x, y*) can be expressed as the product of the phase-noise-free signal and the phase-noise term,

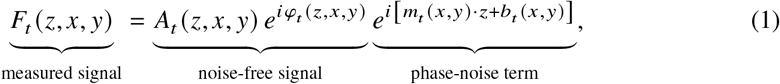

where 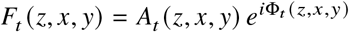 is the complex-valued OCT tomogram with amplitude *A*_*t*_ (*z, x, y*) and phase Φ_*t*_ (*z, x, y*) [29], and *φ*_*t*_ (*z, x, y*) is the noise-free phase. Phase noise can be modeled in most OCT systems as the combination of phase offsets and phase ramps, with corresponding offsets *b*_*t*_ (*x, y*) and slopes *m*_*t*_ (*x, y*) that vary A-line–per–A-line and originate from path-length changes and laser-to-digitizer timing jitter, respectively [30, 31].

OCTA contrast obtained through temporal autocorrelation involves computing complex-conjugate products between measurements at timepoints *t* and *t* + τ, i.e., with time lag τ,

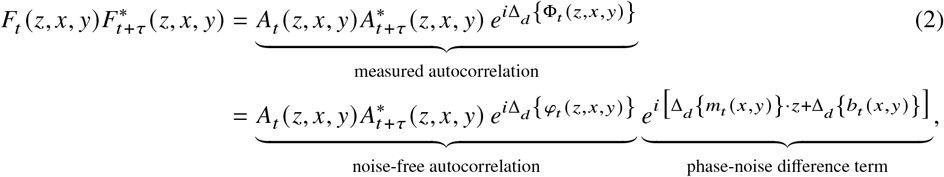

where Δ_*d*_ {(·)} (·)_*t*_ = (·)_*t*_ −(·)_*t* + τ_ are temporal differences and ^∗^ denotes the complex conjugate. The temporal phase-noise difference can be computationally determined and removed with a straight-forward 1D phase-correction in time (Section 3.1); thereby recovering the noise-free complex-conjugate product of interest. This correction leaves the transverse spatial dimensions uncorrected; the key observation is that spatial phase differences do not contribute to the temporal autocorrelation function as demonstrated in Eq. (2). Thus, the phase-corrected complex-conjugate products can be used for 4D ensemble averaging without enforcing phase continuity across spatial dimensions.

### 2.2 Leveraging fully-coherent processing for robust high-contrast continuous-valued 3D OCTA volume calculation

Hereafter, we use *α*_*X*_ to denote the value of an OCTA metric obtained with a given method *X*. As introduced in Section 2.1 (implementation in Section 3.1), we can produce phase-noise-corrected complex-conjugate products which enables us to adapt the first-order autocorrelation function ofthe electric field, 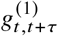, from DLS [32, 33] and its extension to OCT [15, 27],

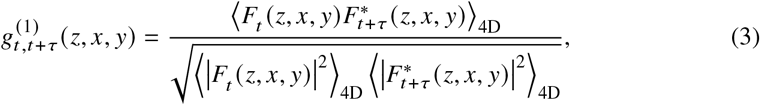

in our coherent OCTA framework with spatial averaging and B-scan repeat protocol, where ⟨ · · · ⟩_4D_ indicates 4D spatiotemporal ensemble averaging. 4D coherent averaging enables high cross-sectional quality (demonstrated in Section 5.4), unlocking the use of 3D vascular metric, which has shown increased disease specificity [22–26], instead of restricting us to 2D metrics based on 2D *en face* projections of retinal layers.

The amplitude of *g*^(1)^ reveals the correlation between repeat measurements which spans a range between fast flow (large vessels, fully decorrelated signal) and “static” (avascular, total signal correlation) tissue, |*g*^(1)^ |∈ [0, 1]. However, it has poor discrimination between static tissue and capillaries, as show in Fig. 1. Resolving the entire 3D microvascular network for detecting capillary dropout and/or angiogenesis is relevant for diseases such as diabetic retinopathy, primary open-angle glaucoma, and age-related macular degeneration [3–6, 34]. This motivates the introduction of the angiographic metric A1C,

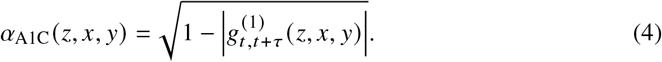

**Fig. 1.**
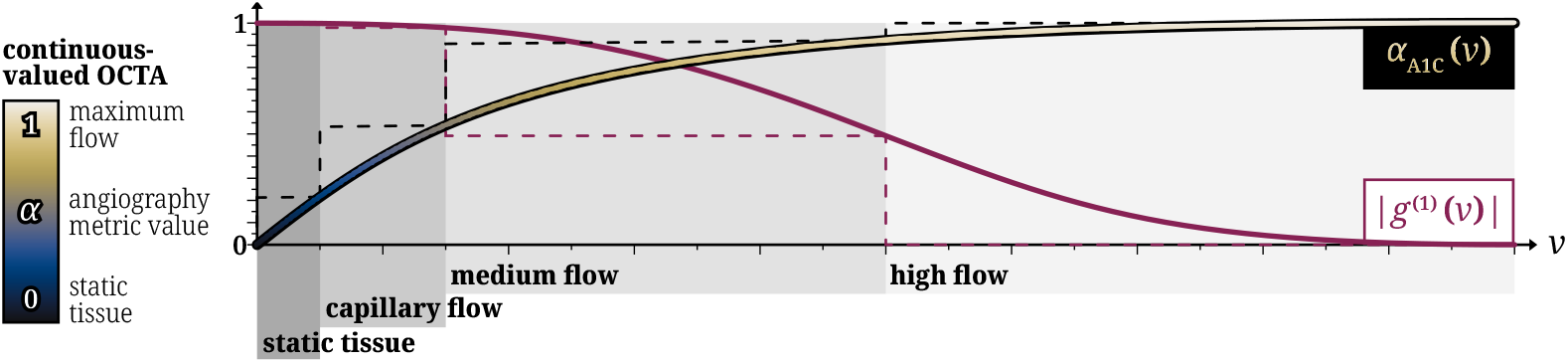
Behavior of |*g*^(1)^ | (solid plum-colored line) and *α*_A1C_ (gradient-colored line) for different flow speeds *v*. Mapping |*g*^(1)^ |into *α*_A1C_ increases the visibility of capillaries against static tissue and matches the convention where zero is no flow and one is maximum flow, while preserving the monotonic relation between flow speed and angiography metric value.

This transformation is adapted from the work by Nam *et al*. [10] to increase capillary delineation and to match the OCTA convention where zero corresponds to no flow and one to maximum flow. Continuous-valued OCTA analysis, sometimes termed “OCTA flux”, has also been found to have improved sensitivity to detect early disease compared with the use of the binarized OCTA signal [20, 21]. Figure 1 shows a comparison between *g*^(1)^ and *α*_A1C_ for various flow speeds using the DLS-OCT model *g*^(1)^ (τ) ∝ exp 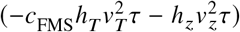, where *h*_*T*_ and *h*_*z*_ are the inverse of the 1/*e* diameter of the transverse and axial point spread functions, respectively, *v*_*T*_ is the transverse speed, *v*_*z*_ is the axial speed [27], and *c*_FMS_ is a correction factor to include effects of forward multiple scattering on the single-scattering DLS model [35]. A1C preserves the monotonic relation between flow speed and |*g*^(1)^ | while offering better quantitative differentiation in the static, slow, and medium speed regimes, beyond binary mapping only.

Additionally, A1C has a closed-form analytical SNR-corrected version, A1C-SNR,

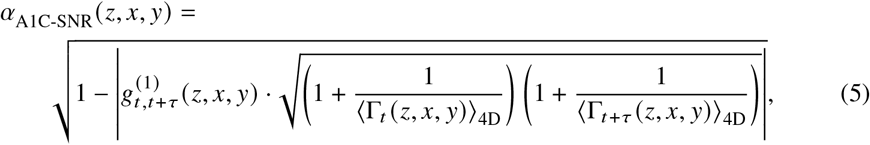

where Γ is the measured SNR value. This expression is derived by modeling the measured OCT signal *F* = *S* + *N* as the composition of the true signal *S* and circular complex-valued Gaussian noise *N* [11, 15] (not to be confused with phase noise). Using the expected value ⟨·⟩ of *N*, ⟨*N*⟩ = 0, and defining SNR as 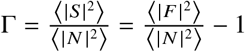, Eq. (5) can be derived from Eq. (4). We will show that, by leveraging 3D/4D ensemble averaging regardless of the amount of phase noise, our estimation of the noisy decorrelation has sufficiently low variance to enable accurate SNR compensation, leading to the recovery of the uncorrupted decorrelation in regions with low SNR and uncovering vessels previously occulted by noise.

## 3. Methods

### 3.1. Temporal phase correction to enable fully-coherent processing

Prior to temporal phase correction, the measured OCT tomograms were corrected for bulk motion between B-scans. Efficient voxel-wise image registration by cross-correlation [36] of the logarithmic-intensity B-scans was used to determine axial and lateral B-scan shifts required to correct for motion between adjacent B-scans. These relative shifts were accumulated to obtain motion correction with respect to the first B-scan, and then high-pass filtered to prevent unintentionally flattening retinal structures. The complex-valued B-scans were padded with the noise floor and then shifted by the accumulated shifts.

Following inter-B-scan bulk motion correction, phase correction was performed along the temporal dimension—across B-scan repetitions—to enable fully-coherent angiography metric calculation. As discussed in Section 2.1, fully-coherent processing for OCTA is enabled with just a 1D phase correction along time using existing techniques such as linear-fit-based [12] or histogram-based [1] methods, even in systems with strong phase jitter such as ours. Retinal tissue locations that are effectively static within the typical OCTA inter-B-scan times of milliseconds are expected to have temporal phase differences values that center around zero in the absence of phase-noise and motion. This also apply to voxels describing purely lateral flow and fully-decorrelating fast flow. Therefore, only voxels in capillaries that have an axial component of flow and are not fully decorrelating should have a non-zero average. Thus, although any phase correction method is suitable, the simpler linear-fit-based 1D phase correction [12] is sufficient to correct for temporal phase-noise between A-lines (see Supplement 1, Section 4 for a diagrammatic demonstration) in this work.

Offsets (*b*_*t*_ *x, y*) and slopes (*m*_*t*_ *x, y*) were determined for every A-line by line fitting the phase differences between temporally adjacent A-lines pair, Δ_*d*_ {Φ_*t*_ (*z, x, y*)}, with weights based on the signal’s logarithmic intensity above the noise floor [12]. To prevent phase wrapping artifacts in the estimation of slopes, the average phase differences ⟨Δ_*d*_ {Φ_*t*_ (*z, x, y*)} ⟩_*z*_ were removed prior to linear fitting; these values were later added to the phase offset found from linear fitting. The calculated phase offsets and phase ramps were then cumulated along the time dimension and applied, such that the A-lines in the first B-scan in the repetition set (*t* = *t*_0_) served as reference for A-lines in subsequent B-scan repetitions (*t* > *t*_0_).

### 3.2. Calculating A1C and A1C-SNR

Section 2.2 defines A1C and A1C-SNR in general form, with time *t* and time lag τ. In the discrete case, the repetition index *r*∈ {0, …, *R*− 1} and step size *d* are defined such that *t* = *r*· Δ*t* and τ = *d*· Δ*t* are multiples of the inter-B-scan time Δ*t*, with *R* total number B-scan repetitions. With the typical step size *d* = 1, we can write

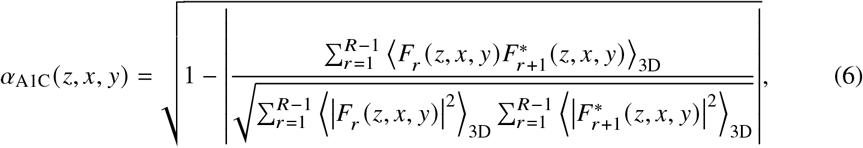

with the temporal averaging made explicit 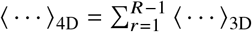. The framework generalizes to any *d* if assessment of decorrelation as a function of τ, such as in capillary flow, is of interest [37]. Because an accurate measure of the SNR is crucial for the correct estimation of the SNR-corrected metric, we assumed that the SNR does not change at a particular voxel as a function of repetition, Γ_*r*_ = Γ_*r*+1_, therefore

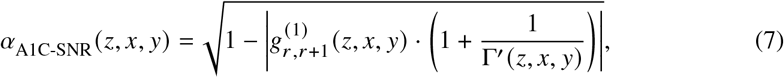

where the SNR for a given spatial location Γ^′^(*z, x, y*) was calculated as

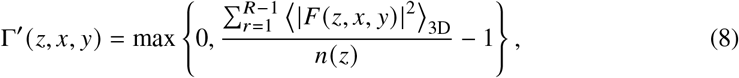

where max {·} is the maximum function, used to avoid negative SNR values, and the depth-dependent noise floor *n* (*z*) was determined as the average signal intensity measured with the light from the sample arm blocked, which is assumed to be constant across A-lines. When appropriate, the SNR in logarithmic scale 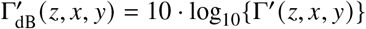 is used.

### 3.3. Characterizing OCTA contrast

To facilitate slicing for analysis and visualization, all volumes were flattened with respect to the retinal pigmented epithelium (RPE) by detecting it and shifting each A-line axially accordingly to the RPE depth.

To effectively evaluate the performance for each OCTA metric, we calculated the contrast-to-noise (CNR) ratio [38], *k*, as

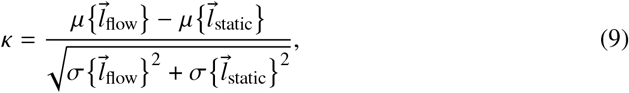

where 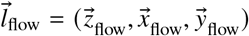 is a region of interest (ROI) belonging to a large blood vessel for which we expect full decorrelation, 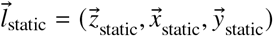 is an ROI belonging to static tissue, and µ{·} and σ{·} represent the mean and standard deviation, respectively. The ROIs were manually selected from a main blood vessel, to capture the largest flow, and from a section of the foveal avascular zone (FAZ) for foveal datasets, and an avascular section of the optic cup for ONH datasets, to capture the background “static” tissue. A larger *κ* indicates a larger separation between the distribution of angiographic values for blood flow and static tissue, quantifying the visibility of vasculature within static tissue. Capillaries with slow flow are more visible in angiography metrics that produce OCTA volumes with greater *κ*. Whereas with decreasing *k*, noise masks low-flow regions, resulting in reduced dynamic range and vasculature delineation.

### 3.4. Facilitating visual comparison with SNR-based masking and range-based rescaling

To suppress voxels with low signal, such those in the vitreous, a mask *M* was applied as 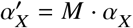 when indicated in the text by “masked”. To generate *M*, a sigmoid function with a growth rate of 2 and an inflection point at 1 was applied to the logarithmic SNR [16],

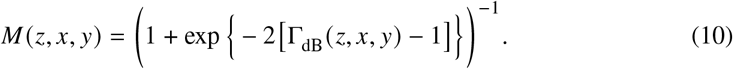

Values *M* < 0.3 were set to zero, effectively discarding voxels with SNR < 0.6 dB with a smooth transition achieved through the sigmoid function.

When indicated by the text “rescaled”, the dynamic range of OCTA visualizations were tuned to enhance visual homogeneity across outputs and allow comparisons between metrics with dataset- or system-dependent scale. The rescaled ranges are given by

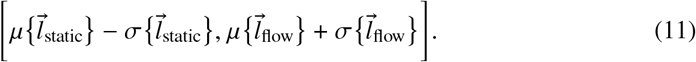

### 3.5. Evaluating depth localization and SNR performance of OCTA volumes

Conventional OCTA visualizations use *en face* projections of the superficial capillary plexus (SCP) and deep capillary plexus (DCP). However, this form collapses most depth information. To capture the 3D localization capabilities of each method, vessel paths from *en face* projections were traced and the resulting cross-sections are displayed (see Supplement 1, Section 6 for details on the projection generation method).

Different retinal layers can have vastly different SNR levels. Traces following layers with different SNRs were also used to create cross-sections to highlight the performance of OCTA metrics at varying SNR levels.

TIFF stacks of the OCTA volumes were exported from MATLAB to ImageJ along with an unsigned 16-bit TIFF stack of 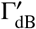. Hand tracing of vessels and depth layers was conducted on TIFF stacks in ImageJ. The freehand line was first converted to a spline through the built-in “fit spline” function and manually fine tuned. Then, the whole TIFF stack was resliced along the spline.

## 4. Imaging Setup

### 4.1. OCT system

We used a custom wavelength-swept OCT system that has been described elsewhere [16]. Briefly, it uses a MEMS-VCSEL laser source (Axsun Technologies, Inc., USA) with a bandwidth of 100 nm centered at 1040 µm and an A-line rate of 100 kHz, a polarization-diverse receiver, and a Signatec PX14400A digitizer. 2048 wavenumber (*k*) samples per A-line were acquired which, after *k*-linearization and dispersion compensation, were zero-padded prior to Fourier transformation to result in A-lines with 2048 depth samples. The laser had a strongly non-linear in *k* sweep, and the Signatec digitizer does not have *k*-clocking capabilities. As a result, the acquired OCT tomograms presented very strong phase noise due to timing jitter between the laser and digitizer, as well as phase noise from axial subject motion and galvanometer scanning. The sample arm was coupled with the ophthalmic interface of a Heidelberg Engineering Spectralis OCT2 system with eye tracking and fixation capabilities, which were used during acquisition. The system had an expected axial resolution with *e*^−2^ diameter of 8.1 µm in air, and the diffraction-limited *e*^−2^ diameter of the point-spread function for a 16.7 mm focal-length eye was 13 µm, considering the double-pass confocal effect.

### 4.2. Imaging protocol

OCT tomograms of a healthy adult volunteer’s fovea and optic nerve head (ONH) were acquired for computing OCTA volumes, and the imaging was conducted in accordance with the tenets of the Declaration of Helsinki [39]. The collected OCT tomograms of the fovea had 4 repetitions per B-scan location, 2048 depth samples, 512 A-lines per B-scan, and 240 B-scan locations (Nyquist sampled in plane, half Nyquist sampled out of plane). This acquisition took 4.92 s and covered a 9.2°× 9.2° FOV (estimated at 5.5 ×5.5 mm^2^ for a 16.7 mm focal-length eye). The collected tomograms of the ONH had 5 repetitions per B-scan location, 2048 depth samples, 512 A-lines per B-scan, and 480 B-scan locations (Nyquist sample both in-plane and out-of-plane). This acquisition took 12.29 s and covered a 9.2° ×9.2° FOV. All OCTA volumes display directly the angiography metric *α* with no intensity masking applied. Only an SNR-based mask is applied when explicitly indicated.

With the inter-B-scan time of ∼5 ms and *e*^−2^ diameter of 13 µm, flow speeds with a lateral component up to 2.6 mm/s will not fully decorrelate *g*^(1)^ and will result in an OCTA flux value, while faster lateral flow speeds will completely decorrelate *g*^(1)^ and will return a plateaued OCTA value. This inter-B-scan time is suitable to return a flux value for typical capillary flows <3 mm/s [37, 40–42], which, although is out of scope for this current study, can support investigations of initial flow alterations at the capillary level.

### 4.3. OCTA processing parameters

In this work, we used our fully-coherent pipeline, FC-OCTA, to generate A1C and A1C-SNR angiography metrics, which are compared against two split-spectrum-based methods, SSADA and SSAPGA, and a partially-coherent method, CDV. Refer to Supplement 1, Section 5, for equations with standardized notation for all methods.

When calculating the split-spectrum methods SSADA and SSAPGA, 11 spectral bins were used with a normalized bandwidth value of 0.28, which have been previously reported as optimal parameters [43]. SSAPGA additionally incorporates an axial phase gradient and its scale factor was set to 2, as previously determined by Liu *et al*. [9]. The normalized bandwidth value of 0.28 has an effective averaging kernel size of 6.2 voxels for our data. To exactly match the amount of averaging among split-spectrum and spatially-averaging methods, the averaging kernel window size, *H*, and *e*^−2^ diameter, *D*, were set to {(*H, D*) : (9, 5.5), (9, 2.7), (5, 1.5)} for {*z, x, y*}, respectively. This 4D axial-centric averaging kernel (temporal and 3D spatial averaging) is the default ensemble averaging kernel used. For the 3D **lateral-only averaging kernel** (temporal and 2D spatial averaging) we used {(*H, D*): (0, 0), (9, 5.6), (5, 2.8)} for {*z, x, y*}, respectively. For both axial-centric and lateral-only averaging kernels the integral of the spatial ensemble equals 6.2 voxels, thus matching the spatial averaging of split-spectrum methods. For *en face* projections in Section 5.4, the 2D ensemble averaging kernel (temporal and 1D spatial averaging) was set to {(*H, D*): (13, 10), (0, 0), (0, 0)} for {*z, x, y*} (with the same 6.2 voxel effective averaging kernel size) to provide only axial averaging prior to layer projection to accomplish fair comparisons with conventional methods.

The *en face* layer projections were based on despeckled intensity tomograms using TNode [44] followed by the segmentation algorithm developed by Mayer *et al*. [45]. See details in Supplement 1, Section 6.

## 5. Results & Discussion

### 5.1. Bias reduction for different averaging approaches

Because of finite sampling, it is only possible to *estimate* the autocorrelation function of the OCT signal. The simulation results presented in Fig. 2 show that the correlation estimation bias (defined in Supplement 1, Section 1), which describes how inaccurate the estimation of the autocorrelation function is, decreases with increasing coherent-to-incoherent averaging ratio. With the increase of temporal averaging (horizontal axis in Fig 2), coherent averaging (A1C) reduces both the mean error and standard deviation, while incoherent averaging (CDV and SSADA) only reduces the standard deviation of the estimation, but not its bias. Interestingly, the errors in the estimated correlation increase with decreasing ground-truth correlation value. This implies that strongly decorrelating regions—blood vessels—benefit the most from the lower bias in fully-coherent ensemble averaging. Overall, A1C presents the lowest bias.

**Fig. 2.**
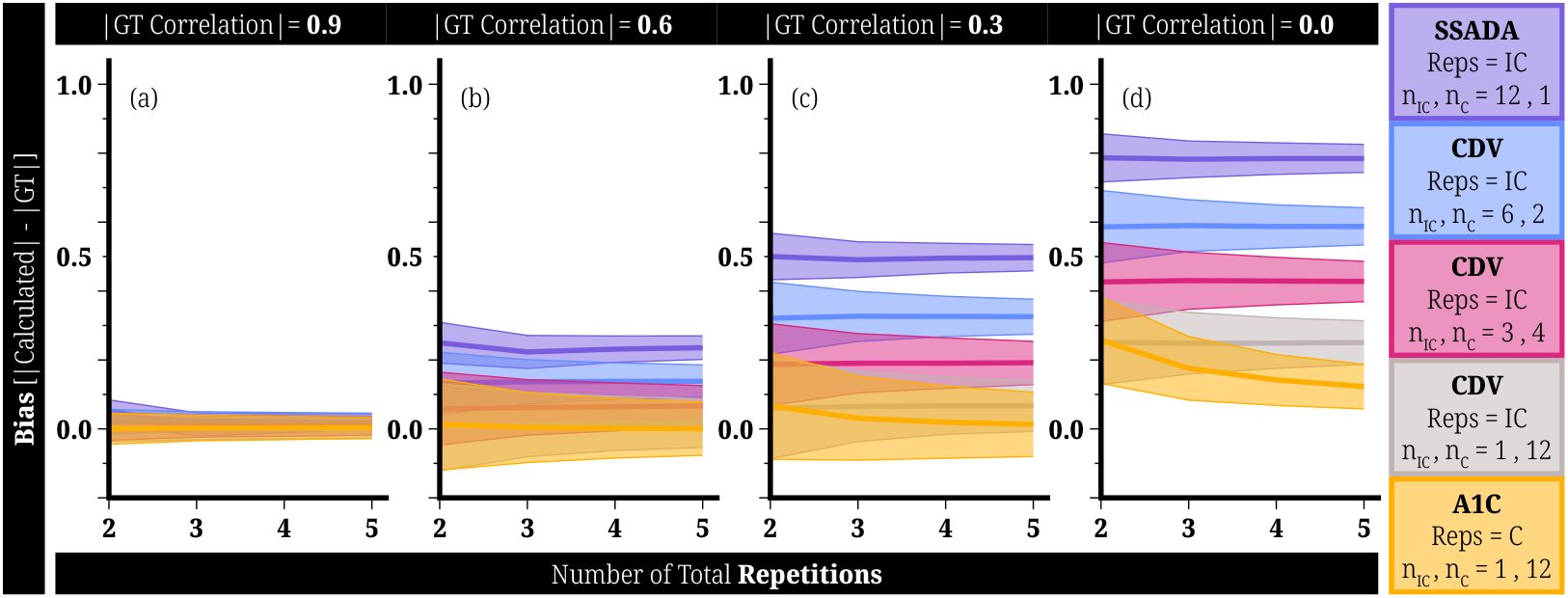
Bias for each simulated combination of coherent and incoherent ensemble averaging plotted versus number of repetitions for various ground-truth (GT) correlation values: (a) high correlation (static tissue), (b) medium correlation, (c) low correlation, and (d) no correlation (fast flow). Fully-coherent A1C consistently produces the lowest estimation bias and is the only method that exhibits a reduction in bias over increasing repetitions. n_C_ ×n_IC_ represents the averaging kernel size in voxels with n_C_ elements for coherent averaging and n_IC_ for incoherent averaging.

Because of this behavior, CDV and SSADA present worse vasculature delineation, lower dynamic range, and CNR compared to A1C and A1C-SNR, as shown in Sections 5.3,5.4, and 5.5.

### 5.2. Temporal phase correction for FC-OCTA

Figure 3 shows that the data acquired in the fovea of a healthy volunteer with our SS-OCT system (without *k*-clock acquisition capabilities) presents significant phase noise prior to temporal phase correction [Fig. 3(b2) shows only the lateral dimension that presents greater phase-noise corruption in the calculation of adjacent A-line phase differences, *y*; see Supplement 1, Section 7 for both lateral dimensions]. Temporal phase differences for static tissue vary in the lateral and temporal dimensions [Fig. 3(b3–b7)]. Direct computation of A1C is therefore impossible [Fig. 3(b)] and only methods that process non-axial dimensions incoherently produce meaningful OCTA volumes. After temporal phase correction (theory in Section 2.1 and implementation in Section 3.1), the distribution of temporal phase differences in static tissue reduces to a narrow Gaussian distribution centered around zero [Fig. 3(f3)], especially after coherent 3D spatial ensemble averaging [Fig. 3(f4)], which is the expected distribution for partially-correlated speckle in absence of deterministic phase shifts, as seen in the phase difference in *z*, where phase is intrinsically stable in Fourier-domain OCT [Fig. 3(b1) and (e1)]. Note that the phase difference between adjacent A-lines in the lateral dimensions remain corrupted after temporal phase correction [Fig. 3(c2) and (f2)] but does not effect the successful removal of temporal phase differences for coherent 4D averaging [Fig. 3(c4–c7) and (f4)]. Thus, temporal phase correction achieves noise-free temporal phase differences across all spatial dimensions for all temporal B-scan pairs [Fig. 3(c4–c7)], enabling FC-OCTA and the computation of A1C and A1C-SNR [Fig. 3(c)].

**Fig. 3.**
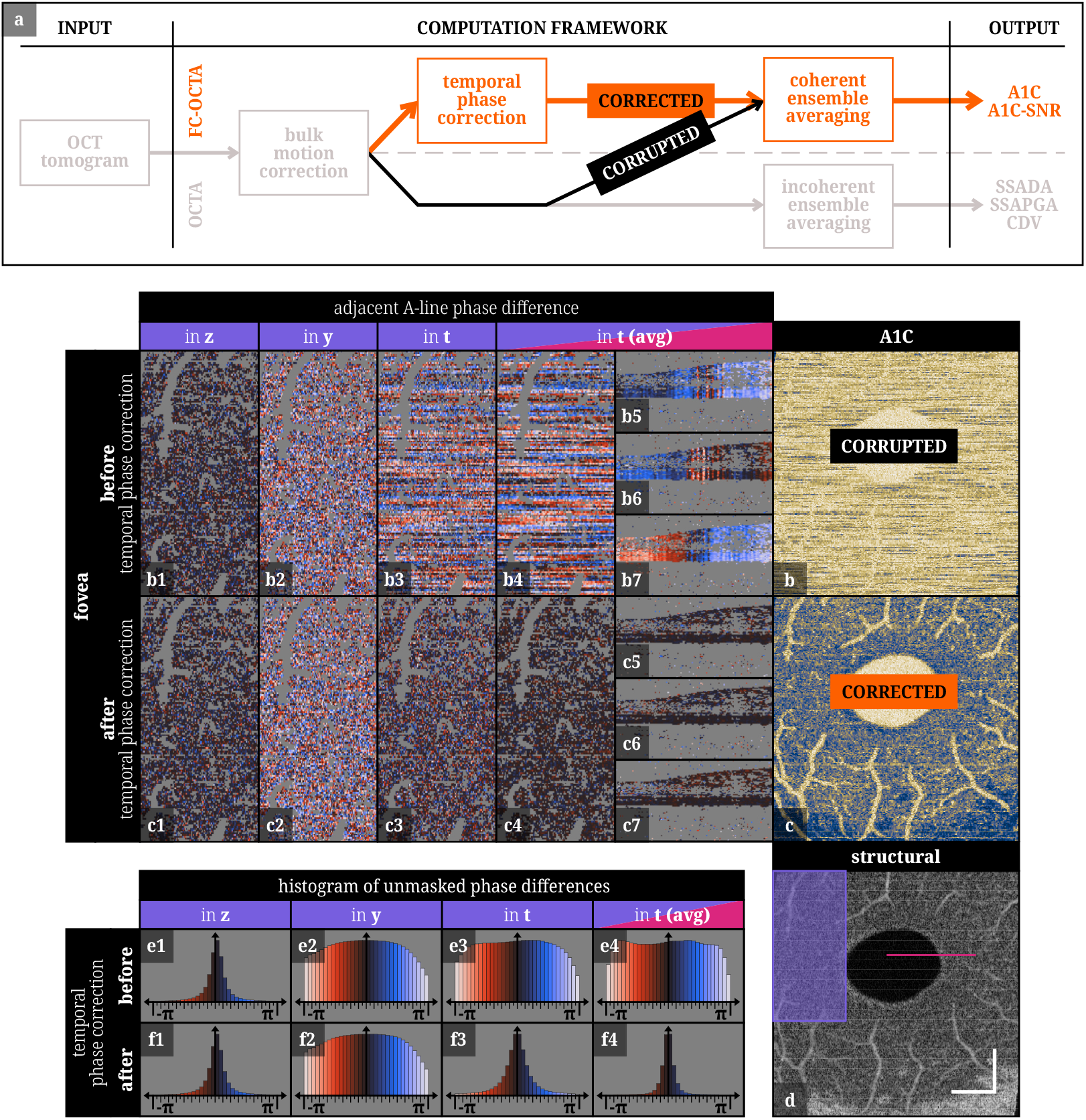
(a) Simplified flow diagram of the processing steps for generating OCTA metrics. FC-OCTA with coherent ensemble averaging is enabled through temporal phase correction; if ignored, (b) the OCTA output volume is corrupted, as opposed to (c) the OCTA output volume is interpretable because the phase was corrected. (d) Two exemplary ROIs of the fovea of a healthy volunteer, purple cropped *en face* and pink cropped B-scan cross-section, are selected to display (b1–b7, c1–c7) the adjacent A-line phase differences before and after temporal phase correction along *z, y, t*, and *t* after coherent 3D spatial ensemble averaging [*t* (avg)] for every temporal B-scan pair (voxels belonging to blood vessels and low-signal regions have been masked out in gray). (e1–e4, f1–f4) Histogram of phase differences were created using the entire volume and all temporal repetitions, ignoring only the grayed-out voxels [(e1) corresponds to (b1), (e2) to (b2), (e3) to (b3), (e4) to (b4–b7), and so forth]. The vertical axis of all histograms has been adjusted for visualization purposes. Scale bars are 1 mm.

### 5.3. OCTA contrast without binarization or projection

We generated OCTA volumes for each angiography metric on both the fovea and ONH datasets. Figure 4 shows *en face* views of the original metrics *α* and after rescaling. We highlight here that these views **correspond to single-depth slabs, not depth-average projections**, an important consideration when interpreting these images. Metrics that use strong axial averaging (SSAPGA, SSADA) will appear to show more/larger vessel segments: this is because many more voxels in depth are merged into a single depth in the final OCTA image. Although this is inconsequential when creating whole-plexus 2D average projections, it is undesirable for 3D analysis of the vascular network.

**Fig. 4.**
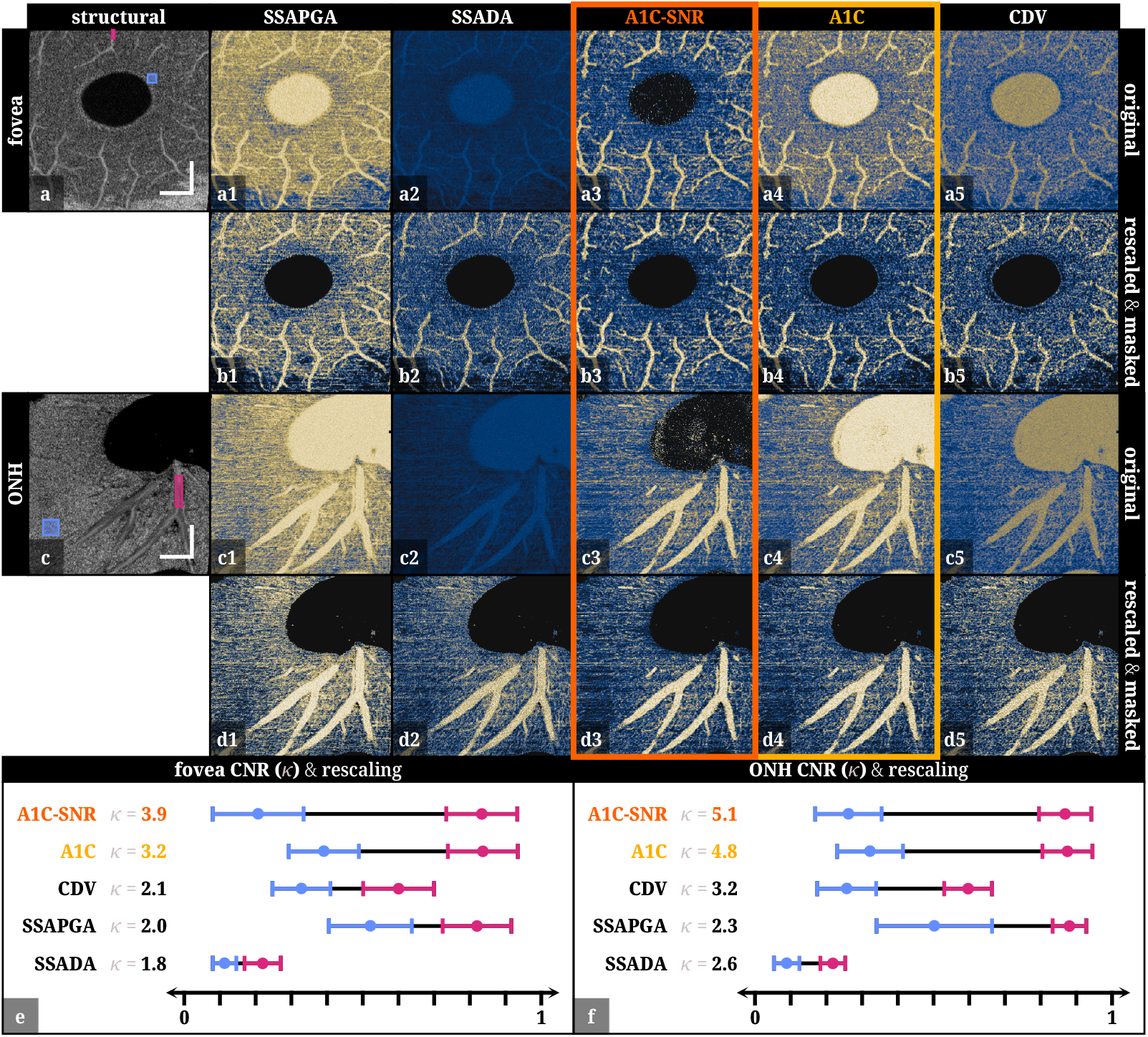
Single-depth *en face* views of OCTA metrics from (a1–a5) the fovea and (c1–c5) optic nerve head of a healthy volunteer, along with (a) and (b) corresponding structural images. Rescaled and masked versions are shown in (b1–b5) for the fovea and (d1–d5) for the optic nerve head. (e) and (f) show the CNR and plots of the mean± standard deviation within the static and blood vessel regions of interest, represented by the blue and pink rectangles in the structural images, respectively. The rescaled limits [Eq. (11)] are indicated by the left end of the blue bar and the right end of the pink bar. A1C-SNR exhibits the largest range and CNR. Scale bars are 1 mm.

The CNR, *k*, for each method was determined using the pink-colored ROI for blood flow and the blue-colored ROI for static tissue showed on the *en face* views displayed on Figs. 4(a) and (c); these ROIs were also used to rescale each OCTA tomogram to aid visual comparison, as discussed in Section 3.4. We also masked out areas with low SNR values as explained in Section 3.4. All subsequent figures present OCTA tomograms with both of these visual transformations.

Consistently across both datasets, *κ* was greatest for A1C-SNR (giving greater than a two-fold increase in contrast from SSADA [Fig. 4(e)] and SSAPGA [Fig. 4(f)]), followed by A1C [Fig. 4(e) and (f)]. This is reflected not only in the greatest separation of the mean *α* value (and *α* distribution) of the blood flow ROI and the static tissue ROI, but also in the dynamic range of the original OCTA volumes, which changed marginally after rescaling. This is manifested in the improved contrast of capillaries in A1C-SNR and A1C, which are less obscured by the high *α* value for static tissue seen in the rest of the metrics [Fig. 4(b1–b5)], and less noisy static tissue values at regions selected to produce the best possible *κ* [Fig. 4(d1–d5)]. Visualizations 1 and 2 contain fly-through views for all rescaled and masked metrics for the fovea and ONH datasets, respectively, enabling a detailed comparison of the image quality. Alternatively, refer to Supplement 1, Section 8 for a zoomed-in ROI version of Fig. 4.

From our results, it is clear that A1C and A1C-SNR provide OCTA images with the best microvasculature delineation, while preserving the quantitative nature of *α* and without relying on rescaling in the case of A1C-SNR.

The increase in contrast performance by A1C-SNR is better appreciated at a deeper *en face* view with lower SNR (Fig. 5). Figure 5 shows a zoomed-in ROI [Fig. 4(c) and (c1–c5)] with two circled regions that highlight foveal capillary paths only confidently identifiable amongst the background static tissue with A1C-SNR [Fig. 5(c3)]. A more detailed contrast performance analysis across SNR levels of OCTA metrics is provided in Section 5.5.

**Fig. 5.**
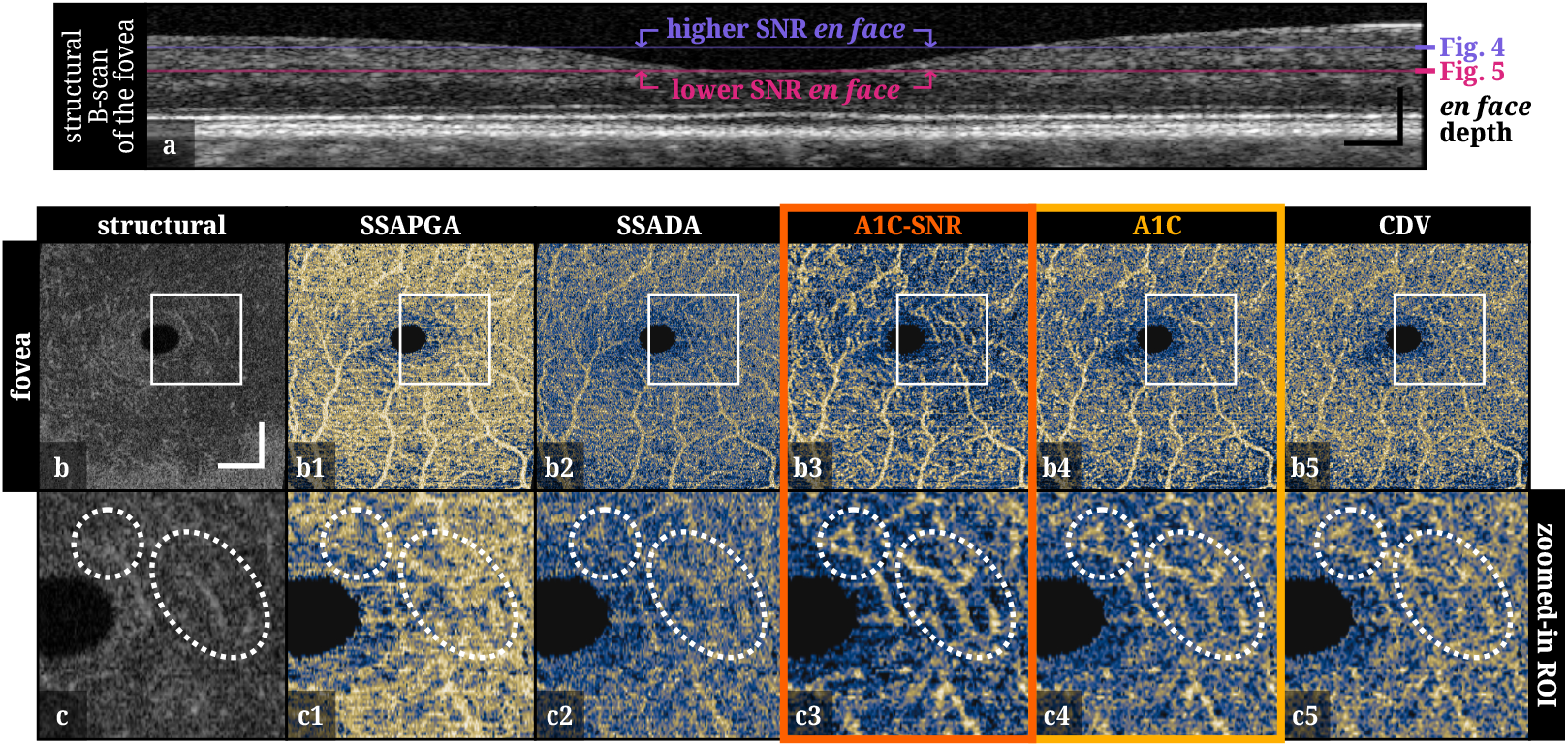
(a) Exemplary structural B-scan cross-section of the fovea shows the depth locations of the selected single-depth *en face* views of Fig. 4 and Fig. 5. (b1–b5) Single-depth *en face* view of the OCTA metrics with (b) the corresponding structural image. The white rectangular area is zoomed-in to for (c1–c5) the OCTA metrics and (c) the corresponding structural image. The dotted white circle and ellipse ROIs feature capillary paths that are recovered through A1C-SNR. White and black scale bars are 1 mm and 0.25 mm, respectively.

### 5.4. Depth localization of microvasculature

Figure 6 shows 75^th^ percentile projections of the SCP and DCP plexuses for masked and rescaled OCTA volume of the fovea dataset from Fig. 4. Clearly, the CNR benefits of coherent processing also translate to projection images, with capillaries being significantly more visible in A1C-SNR. Binarization and vessel segmentation is typically performed on these projection images to calculate diagnostic metrics such as vessel area density, branchpoint density, and vessel tortuosity [18]. It is expected that the improvement of contrast of the vasculature would improve the performance of further analysis protocols.

**Fig. 6.**
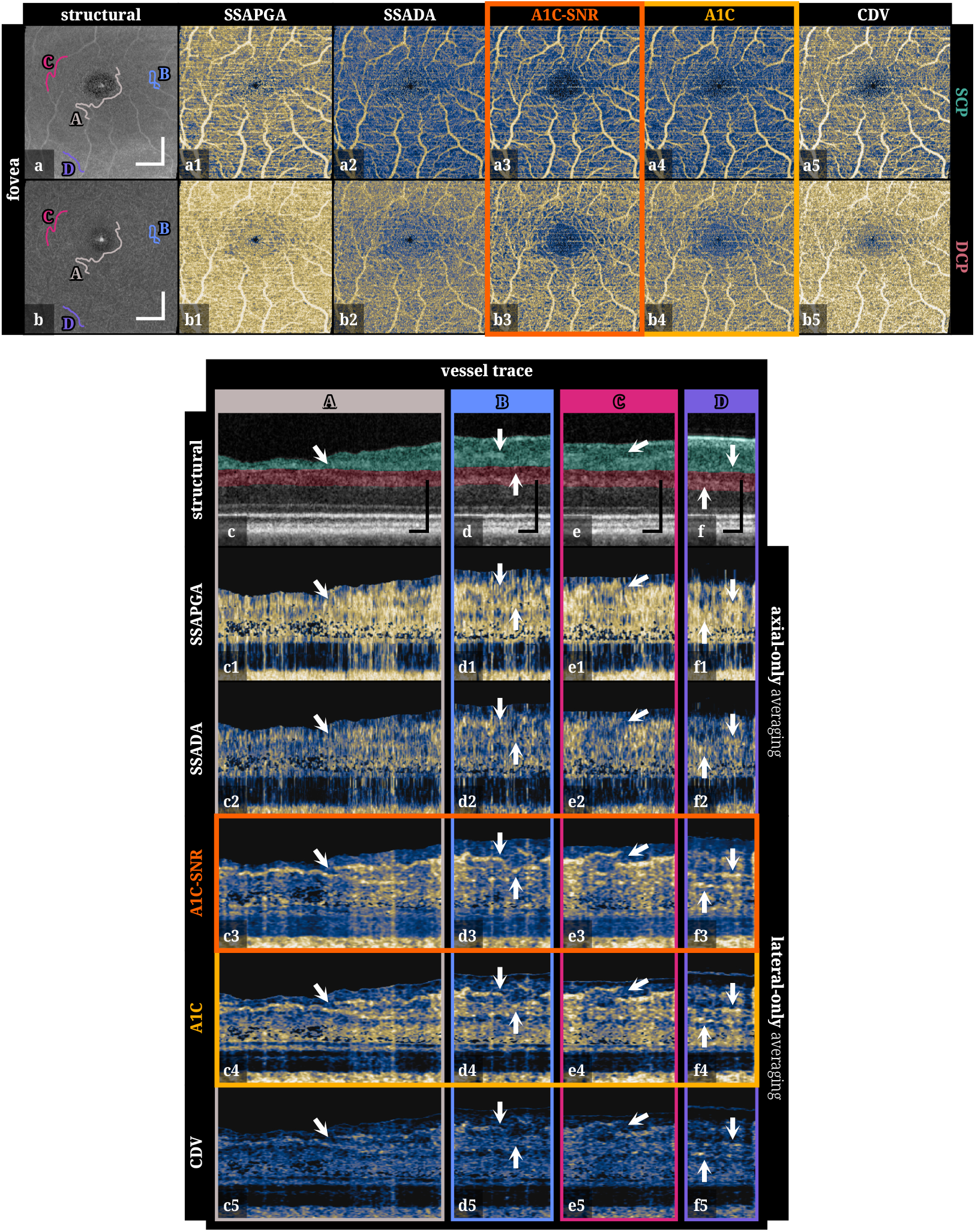
75^th^ percentile projections of the capillary plexuses, (a1–a5) SCP and (b1–b5) DCP, performed on the rescaled OCTA volumes for each angiography metric. (a) and (b) structural projections showing four vessel traces used to generate cross-sectional views with distinct features: (A) overlapping blood vessels, (B) sharp change in depth, (C) disconnected blood vessels, and (D) blood vessels in two distinct layers. White and black scale bars are 1 mm and 0.25 mm, respectively.

However, as mentioned earlier, studies have found that 3D version of diagnostic metrics can be more specific for disease identification and staging [22–26], in which case projection visualization are not suitable. Although SSADA, SSAPGA, and CDV are usable for projection analyses [Fig. 6(a1–a5)], they struggle in providing interpretable depth localization of microvasculature [Fig. 6(c1–c5), (d1–d5), (e1–e5), and (f1–f5)]. FC-OCTA provides versatility to shape the spatial ensemble average in A1C in any way, opening the door for averaging strategies that are more suitable for 3D OCTA post-processing than the axial averaging that is predominantly used. To further explore the implications of 3D OCTA, we examined and traced vessels from the projection images [Fig. 6(c), (d), (e), and (f)]. The difference in vessel contrast in cross-sectional images is striking among the different angiography metrics, with particularly A1C-SNR delivering excellent cross-sectional contrast which preserves depth resolution. These cross-sectional vessel traces exhibit separate vessels that clearly lie at different depths but overlap in *en face* projections. The ability to preserve depth information in 3D vascular metrics is critical in these instances and may be crucial in why these metrics are more specific than 2D vascular metrics.

Ensemble averaging methods posses flexibility to leverage different amounts of axial and lateral averaging, unlike split-spectrum methods (SSADA and SSAPGA), which are limited to the axial dimension. A1C, A1C-SNR, and CDV are able to leverage lateral-only averaging (discussed in Section 4.3) to increase depth clarity, and A1C and A1C-SNR have coherent averaging further maximizing the efficiency of averaging (CDV uses incoherent averaging for non-axial dimensions).

Thus, A1C-SNR and A1C not only increase contrast of OCTA volumes and maintain its quantitative *α* value, but also improve on the preservation of volumetric information. These advantages of FC-OCTA make this method more effective at producing 3D microvasculature maps and potentially more effective at creating inputs for further 3D diagnostic analyses.

### 5.5. Visibility across SNR levels

Across OCT tomograms, regions vary in SNR due to varying scattering properties of tissue and signal attenuation with depth. In the retina, the deeper transparent layers (inner nuclear layer, outer nuclear layer) have lower SNR, which negatively impacts the contrast of angiography metrics because decorrelation from noise becomes a significant contributor to the OCTA signal.

To further study the effect of SNR in OCTA, Fig. 7 shows four different depths that were hand-traced in the ONH dataset, along with ROIs selected within these depth sections to illustrate their average 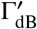 value. A1C-SNR implements SNR correction on A1C (theory Section 2.2, implementation Section 3.2) and is able to recover capillary contrast across all three low SNR scenarios [Fig. 7(b3), (c3), (d3), and (e3)].

**Fig. 7.**
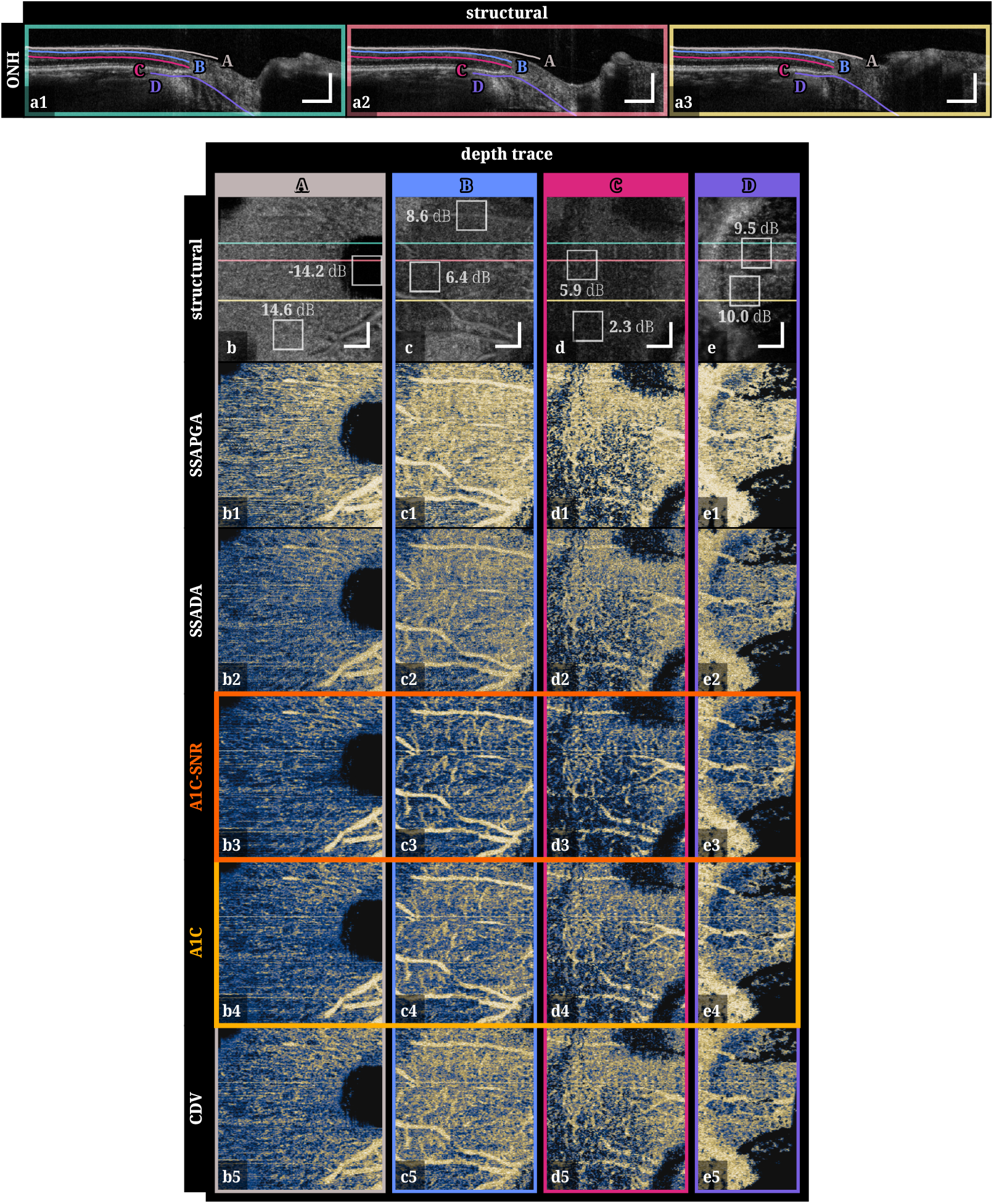
*En face* views of the optic nerve head following the traces shown in the B-scans in (a1–a3) with various SNR levels: (A) nerve fiber layer (high SNR), (B) ganglion cells layer (lower SNR), (C) inner nuclear layer (even lower SNR), and (D) underneath the retinal pigmented epithelium and into the lamina cribrosa. The structural *en face* images in (b–e) include the mean SNR within the white boxes and colored horizontal lines representing the location of the B-scans. Scale bars are 0.5 mm.

Depth trace D, which selects the depth underneath the RPE that enters the laminar region of the ONH, is particularly compelling as most angiography exploration for diagnostics have been performed on the papillary and peripapillary vessels that surround the optic disc [34, 46–48]. Laminar region analyses have been mostly confined to structural imaging [49]. However, changes in these regions have been linked to myopia and glaucoma development [50]. Therefore, enabling the investigation of capillaries at this depth with A1C-SNR may provide further diagnostic material for diseases impacting the ONH.

## 6. Conclusion

In this work, we demonstrated the possibility of using complex-valued OCT tomograms acquired through conventional raster scanning for OCTA calculation regardless of the presence of system phase instability. We presented a method for computing OCTA, FC-OCTA, which includes a new fully-coherent OCTA metric, A1C, that includes a closed-form analytical SNR correction, A1C-SNR, and a computational framework that resolves any prior system phase-noise for autocorrelation analysis. Although phase noise can only be robustly corrected along a single dimension at a time, we demonstrated this is sufficient even when coherent 4D spatio-temporal ensemble averages are used.

Previous methods of OCTA calculation retain one or more dimensions of incoherent averaging, which results in their reduced CNR from hindered bias reduction capabilities, ultimately reducing delineation of microvasculature. We conducted an analysis of strategies for bias reduction in OCTA and demonstrated coherent averaging, especially along the temporal dimension, is crucial to minimizing estimation bias. Beyond showing improved contrast with FC-OCTA, we demonstrated kernel shaping flexibility with A1C and A1C-SNR, enabling better depth localization of capillary paths and the visibility of 3D features such as overlapping vessels. We also showed that our SNR-corrected metric, A1C-SNR, is able to, in a system with strong phase noise, resolve capillaries in much deeper and lower SNR retinal layers that was not identifiable with other OCTA methods, in addition to providing the best overall contrast. We believe that these attributes will enable high-fidelity 3D diagnostic analyses and OCTA investigation of deeper regions of the ONH.

To the best of our knowledge, FC-OCTA with A1C and A1C-SNR is the first fully-coherent and phase-noise-tolerant method for OCTA. All together, FC-OCTA provides a robust method to construct volumetric OCTA that we envision will improve the viability of clinical adoption of OCTA in addition to its diagnostic sensitivity.

## Data Availability Statement

Data may be obtained from the authors upon reasonable request. Sample processing and test data are available at Code 1, Ref. [51] and Dataset 1, Ref. [52].

## Supplemental document

See Supplement 1 for supporting content.

