## Supplemental Document for "Phase-noise tolerant fully-coherent optical coherence tomography angiography for high-contrast 3D microvasculature mapping"

### 1. DEMONSTRATING REDUCED STATISTICAL BIAS WITH 4D COHERENT ENSEMBLE AVERAGING

To evaluate the performance of an angiography metric in autocorrelation estimation and bias reduction, we implemented numerical simulations to investigate the error between the ground-truth and estimated correlation values for repeated measurements. The equations for the various angiography metrics discussed in this paper are provided with standardized notation in Section 5. Hereafter, following conventional nomenclature, we denote the numerator of an autocorrelation metric (that is, the "unnormalized" autocorrelation) as the covariance. Each angiography metric has different definitions of the autocovariance (in particular, the ensemble averaging methodology), and its normalization, $\eta_X\{\cdots\}$. We define the autocorrelation of the OCT signal for method $X$ as $\beta_X$. For A1C, $\alpha_{\text{A1C}} = \sqrt{1-\beta_{\text{A1C}}}$. In Section 2, we demonstrate that the effect of $\eta_X\{\cdots\}$ is minimal in terms of statistical bias, so we focus the analysis here on the contribution of coherent averaging towards bias reduction.

The simulation framework generated a temporal series $\{m_i : i = 1, 2, ..., R\}$ with $R$ observations using the recurrence relation

$$m_i = C \cdot m_{i-1} + \sqrt{1-C^2} \cdot s_i, \tag{S1}$$

where $m_i$ represents the observation at time-point $i$. We generated $L$ independent trials of each temporal series to analyze its statistical behavior. $C \in [-1, 1]$ is the ground-truth correlation, $\{s_i : i = 1, 2, \cdots, R\}$ are independent zero-mean circular Gaussian realizations, and the base case of the recursion is $m_1 = s_1$. For all $L$ independent trials, $s_i$ was drawn from the same complex, circular Gaussian distribution. See Section 3, for the theoretical derivation of the recurrence relation. This procedure was performed in parallel for a 2D grid of $M \times M$ distinct spatial locations to allow for coherent and incoherent ensemble averaging along orthogonal dimensions with kernel sizes $n_{\text{C}}$ and $n_{\text{IC}}$, respectively, effectively emulating various ratios of coherent-to-incoherent averaging (see Fig. S1).

The estimated correlation value with a given metric $X$ was calculated with either

$$\beta_X = \eta_X \left\{ \left\langle \left| \langle m_i \cdot m_{i+1}^* \rangle_{n_{\text{C}},R} \right| \right\rangle_{n_{\text{IC}}} \right\} \quad \text{or} \quad \beta_X = \eta_X \left\{ \left\langle \left| \langle m_i \cdot m_{i+1}^* \rangle_{n_{\text{C}}} \right| \right\rangle_{n_{\text{IC}},R} \right\}, \tag{S2}$$

depending on how each metric implements temporal averaging of repetitions (coherent or incoherent averaging). The error between the ground-truth and estimated correlation values for each angiography metric, $\varepsilon_X$, was calculated as

$$\varepsilon_X = \beta_X - |C|, \tag{S3}$$

from which we computed the mean and standard deviation cross all $L$ trials to quantitatively compare the correlation estimation bias of different metrics.

#### A. Simulation processing parameters

To compare the reduction of bias of different ensemble averaging strategies when estimating autocorrelation during OCTA, we used $\beta_{\text{SSADA}}$ for fully-incoherent processing, $\beta_{\text{CDV}}$ for partially-coherent processing, and $\beta_{\text{A1C}}$ for fully-coherent processing. Following the simulation framework

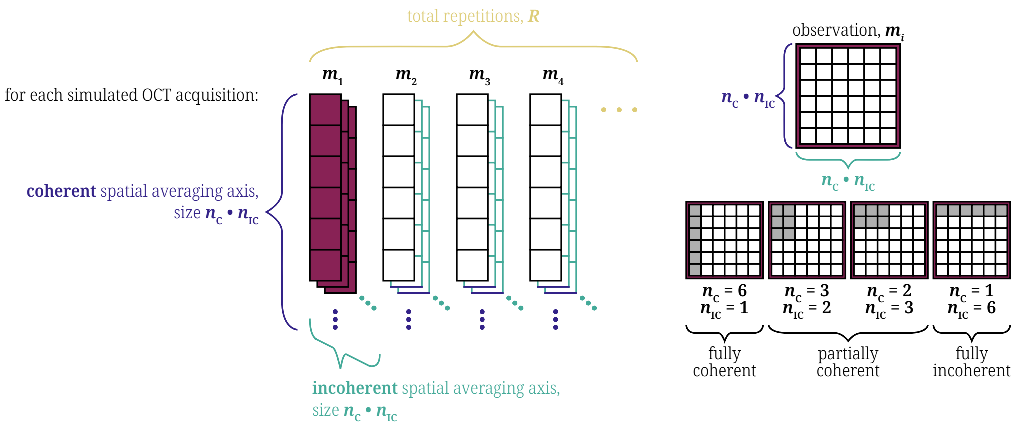


**Fig. S1.** For each simulated OCT acquisition with $R$ repetitions, each $m_i$ had a square grid with side of length $M = n_C \cdot n_{IC}$. To test various contributions of coherent and incoherent averaging, rectangular sub-grids were selected such that every variation contains the same amount of voxels. The vertical axis was used for coherent spatial averaging with kernel size $n_C$, while the horizontal axis was used for incoherent spatial averaging with kernel size $n_{IC}$. Finally, averaging across $m_i \cdot m_{i+1}^*$ was chosen to be completed either coherently or incoherently. C: coherent ensemble averaging, IC: incoherent ensemble averaging.

discussed in Section 1, we simulated $R = 5$ observations with $L = 1 \times 10^4$ independent trials and $M = 12$ spatial locations. Defining $Q$ as the binary choice of whether to coherently average ($Q = 1$) or incoherently average ($Q = 0$) the temporal dimension in the calculation of the estimated correlation value $\beta_X$, the set of coherent-to-incoherent averaging combinations tested was $\{(n_C, n_{IC}, Q) : (12, 1, 1), (12, 1, 0), (4, 3, 0), (2, 6, 0), (1, 12, 0)\}$. **Results of this simulation is presented in the main article, Section 5.1.**

### 2. NORMALIZATION METHOD HAS MINIMAL INFLUENCE ON STATISTICAL BIAS

In estimating the autocorrelation, metrics divide out the mean to normalize the correlation value, $\beta_X = \eta_X\{\cdots\}$. There are multiple ways of computing the mean $\mu\{\cdots\}$: root-mean-squared mean (RMS) $\mu_{\text{RMS}}\{\cdots\}$, arithmetic mean (AM) $\mu_{\text{AM}}\{\cdots\}$, geometric mean (GM) $\mu_{\text{GM}}\{\cdots\}$, and harmonic mean (HM) $\mu_{\text{HM}}\{\cdots\}$,

$$\underbrace{\sqrt{\frac{I_1^2 + I_2^2}{2}}}_{\text{RMS}} \geq \underbrace{\frac{I_1 + I_2}{2}}_{\text{AM}} \geq \underbrace{\sqrt{I_1 I_2}}_{\text{GM}} \geq \underbrace{\frac{2}{\frac{1}{I_1} + \frac{1}{I_2}}}_{\text{HM}}, \tag{S4}$$

respectively. The normalization method in $g^{(1)}$ (and therefore A1C) is GM $\eta_{\text{A1C}}\{\cdots\} = \frac{\cdots}{\mu_{\text{GM}}\{\cdots\}}$, whereas in SSADA and CDV it is AM $\eta_{\text{A1C}}\{\cdots\} = \frac{\cdots}{\mu_{\text{AM}}\{\cdots\}}$. To compare only the effect of the normalization method on bias reduction, maintaining a fully coherent ensemble averaging across all cases, A1C's denominator was modified to evaluate each of the four types of normalization: A1C-RMS, A1C-AM, A1C-GM, and A1C-HM (see Fig. S2). No significant contrast difference was observed both qualitatively and quantitatively evaluating the CNR (method discussed in main article, Section 3.3) for the four normalization methods.

Based on these results, we chose to use the GM version as it follows the standard definition of the first-order autocorrelation function and it provides a straightforward analytical SNR correction [main article, Eq. (5)].

### 3. THEORETICAL DERIVATION OF MULTIPLE PARTIALLY CORRELATED OCT SIGNALS TO SIMULATE B-SCAN REPETITIONS

#### A. Case of two repetitions

For two uncorrelated OCT signals arising from the same complex-valued Gaussian distribution, $s_1$ and $s_2$, the expected values satisfy $\langle |s_1|^2 \rangle = \langle |s_2|^2 \rangle = I$ and $\langle s_1 s_2^* \rangle = \langle s_1^* s_2 \rangle = 0$. Considering

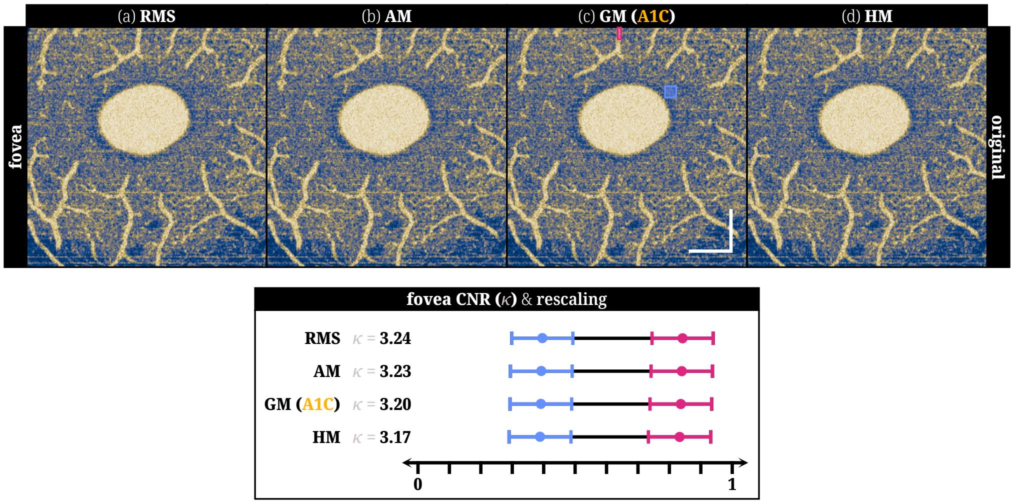


**Fig. S2.** Four modified A1C metrics were computed using the fovea dataset described in the main article, Section 4.2. Each of the modified A1C metrics used one of the four normalization methods for the autocorrelation function denominator: (a) A1C-RMS, (b) A1C-AM, (c) A1C-GM (the normal A1C metric), and (d) A1C-HM. We observed that the CNR values follow the same relation in Eq. (S4). However, the contrast differences are **two orders of magnitude smaller** than the CNR differences between different angiography metrics (seen in main article, Fig. 4). Therefore, the mean of normalization (whether taking RMS, AM, GM, or HM) has no significant impact on the inherent statistical bias a given angiography metric provides.

a first measurement $m_1 = s_1$, a partially correlated second measurement has the form

$$m_2 = c_1 m_1 + c_2 s_2, \tag{S5}$$

where $c_1$ and $c_2$ are scalar factors that we will define. Because we consider that the intensity of the OCT signal is constant in time, the two measurements come from the same distribution and $\langle |m_1|^2 \rangle = \langle |m_2|^2 \rangle = I$. If the two measurements have a correlation value $C$, then the first-order autocorrelation function becomes

$$C = \frac{\langle m_1 m_2^* \rangle}{\sqrt{\langle |m_1|^2 \rangle \langle |m_2|^2 \rangle}}. \tag{S6}$$

Therefore, Eqs. (S6) and (S5) give

$$C = \frac{c_1^* I + 0}{\sqrt{I^2}} \tag{S7}$$

$$I = |c_1|^2 I + 0 + 0 + |c_2|^2 I \tag{S8}$$

so $c_1 = C^*$ and $c_2 = \sqrt{1 - |C|^2}$. Given that we always take the absolute value of the correlation value $C$ to calculate any of the OCTA metrics considered in this work, we can limit our investigation to non-negative real-valued correlations and define $C \in \mathbb{R}_0^+$,

$$m_2 = C \cdot m_1 + \sqrt{1 - C^2} \cdot s_2. \tag{S9}$$

### B. Case of multiple repetitions

In this case, each new measurement has a correlation value of $C$ with respect to the prior measurement. The third measurement $m_3$ would then be related to $m_2$ by

$$m_3 = c_1 m_2 + c_2 s_3, \tag{S10}$$

where $\{c_1, c_2\}$ are a new set of some scalar factors. Again, we define $\langle |m_3|^2 \rangle = I$ and since $s_3$ is a totally uncorrelated signal to $s_1$ and $s_2$, $\langle m_2 s_3^* \rangle = \langle m_2^* s_3 \rangle = 0$. This pair of measurements will still also be correlated by $C$, so

$$C = \frac{\langle m_2 m_3^* \rangle}{\sqrt{\langle |m_2|^2 \rangle \langle |m_3|^2 \rangle}}. \tag{S11}$$

Using Eqs. (S5) and (S11), we once again find that $c_1 = C^*$ and $c_2 = \sqrt{1 - |C|^2}$, and after similarly limiting our investigation to $C \in \mathbb{R}_0^+$, $m_3 = C \cdot m_2 + \sqrt{1 - C^2} \cdot s_3$. Therefore, the case of multiple repetitions is an iterative generalization of the two-repetition case, with the base case $m_1 = s_1$ and the iterative step

$$m_i = C \cdot m_{i-1} + \sqrt{1 - C^2} \cdot s_i. \tag{S12}$$

### 4. PREDOMINANT ZERO-MEAN TEMPORAL PHASE DIFFERENCE ALLOWS FOR STRAIGHT-FORWARD TEMPORAL PHASE CORRECTION

As mentioned in the main article, Section 3.1, retinal tissue locations that are effectively static and blood vessel locations that fully decorrelate or demonstrate purely lateral flow all are expected to have temporal phase difference values that center around zero in the absence of phase noise and motion. Therefore, the simpler linear-fit-based 1D phase correction [1] is sufficient to correct for temporal phase noise between A-lines (Fig. S3).

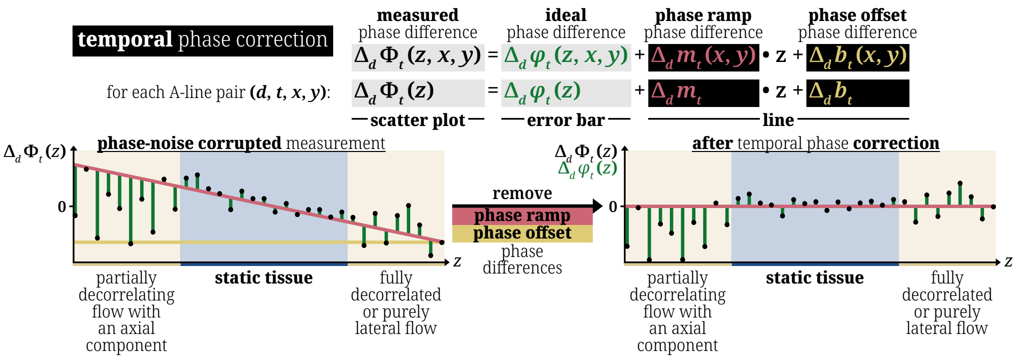


**Fig. S3.** Temporal phase correction finds and removes the linear fit of the phase difference between every temporal A-line pair.

### 5. COMPARISON WITH OTHER ANGIOGRAPHY METRICS

Here, we write the equations for all angiography metrics used in this work with standardized notation matching main article, Eq. (6) for ease of comparison and replication. Every angiography metric has a range of [0, 1].

#### A. SSADA [2]

$$\alpha_{\text{SSADA}}(z,x,y) = 1 - \beta_{\text{SSADA}}(z,x,y) = 1 - \left\langle \frac{2\,|F_{w,r}(z,x,y)|\,|F_{w,r+1}(z,x,y)|}{|F_{w,r}(z,x,y)|^2 + |F_{w,r+1}(z,x,y)|^2} \right\rangle_{W,R} \tag{S13}$$

where $w$ is the spectrum window in one of the total $W$ binning windows taken. It was shown that $W = 11$ with a normalized bandwidth value of 0.28 gave the best results [3] and that was used in our studies for SSADA and SSAPGA.

#### B. SSAPGA [4]

$$\alpha_{\text{SSAPGA}}(z,x,y) = 1 - \beta_{\text{SSAPGA}}(z,x,y) = 1 - \left\langle \frac{2\,|F_{w,r}(z,x,y)|\,|F_{w,r+1}(z,x,y)|\,\left| e^{j\rho \frac{\mathrm{d}(\angle[F_{w,r+1}(z,x,y)] - \angle[F_{w,r}(z,x,y)])}{\mathrm{d}z}} \right|}{|F_{w,r}(z,x,y)|^2 + |F_{w,r+1}(z,x,y)|^2} \right\rangle_{W,R} \tag{S14}$$

where $\frac{\mathrm{d}(\angle[F_{w,r+1}(z,x,y)]-\angle[F_{w,r}(z,x,y)])}{\mathrm{d}z}$ is the axial phase gradient and $\rho = 2$ is the optimized scale factor found by Liu *et. al.* and it is the value we used in our studies [4].

#### C. CDV [5]

$$\alpha_{\mathrm{CDV}}(z,x,y) = \sqrt{1-\beta_{\mathrm{CDV}}(z,x,y)} = \sqrt{1-\frac{2\left\langle\left|\left\langle F_r(z,x,y)F^*_{r+1}(z,x,y)\right\rangle_{\mathrm{axial}}\right|\right\rangle_{\mathrm{lateral},R}}{\left\langle|F_r(z,x,y)|^2+|F_{r+1}(z,x,y)|^2\right\rangle_{4D}}} \tag{S15}$$

where $\langle\cdots\rangle_{\mathrm{axial}}$ is coherent 1D axial averaging and $\langle|\cdots|\rangle_{\mathrm{lateral}}$ is incoherent 2D lateral averaging.

### 6. METHOD TO GENERATE SCP AND DCP *EN FACE* PROJECTIONS

The raw OCT tomogram was first despeckled using TNode [6]. A retinal segmentation algorithm based on the work by Mayer *et al.* [7] was used on the despeckled tomogram to produce a matrix labeling the layer interfaces between the vitreous fluid, retinal nerve fiber layer (RNFL), ganglion cell layer (GCL), inner plexiform layer (IPL), inner nuclear layer (INL), outer plexiform layer (OPL), and outer nuclear layer (ONL). The superficial capillary plexus (SCP) binary mask was created using the volume between the vitreous-fluid-to-RNFL interface to the IPL-to-INL interface. The deep capillary plexus (DCP) binary mask was created using the volume between the IPL-to-INL interface to the OPL-to-ONL interface. These masks were applied to the SNR-masked and rescaled OCTA volumes before a given percentile *en face* projection was calculated.

### 7. FIGURE 3 WITH ALL SPATIOTEMPORAL DIMENSIONS

In Figure 3 of the main article, we only show the lateral dimension that presents greater phase-noise corruption in the calculation of adjacent A-line phase differences, $y$. Figure S4 is Fig. 3 but with both lateral dimensions, $x$ and $y$, to illustrate the behavior of all spatiotemporal dimensions after temporal phase correction.

### 8. ZOOMED-IN FIGURE 4

In Figure 4 of the main article, we show the full single-depth *en face* view. For Figure S5, we provide a magnified view of each *en face* image to aid the visual comparison of each metric.

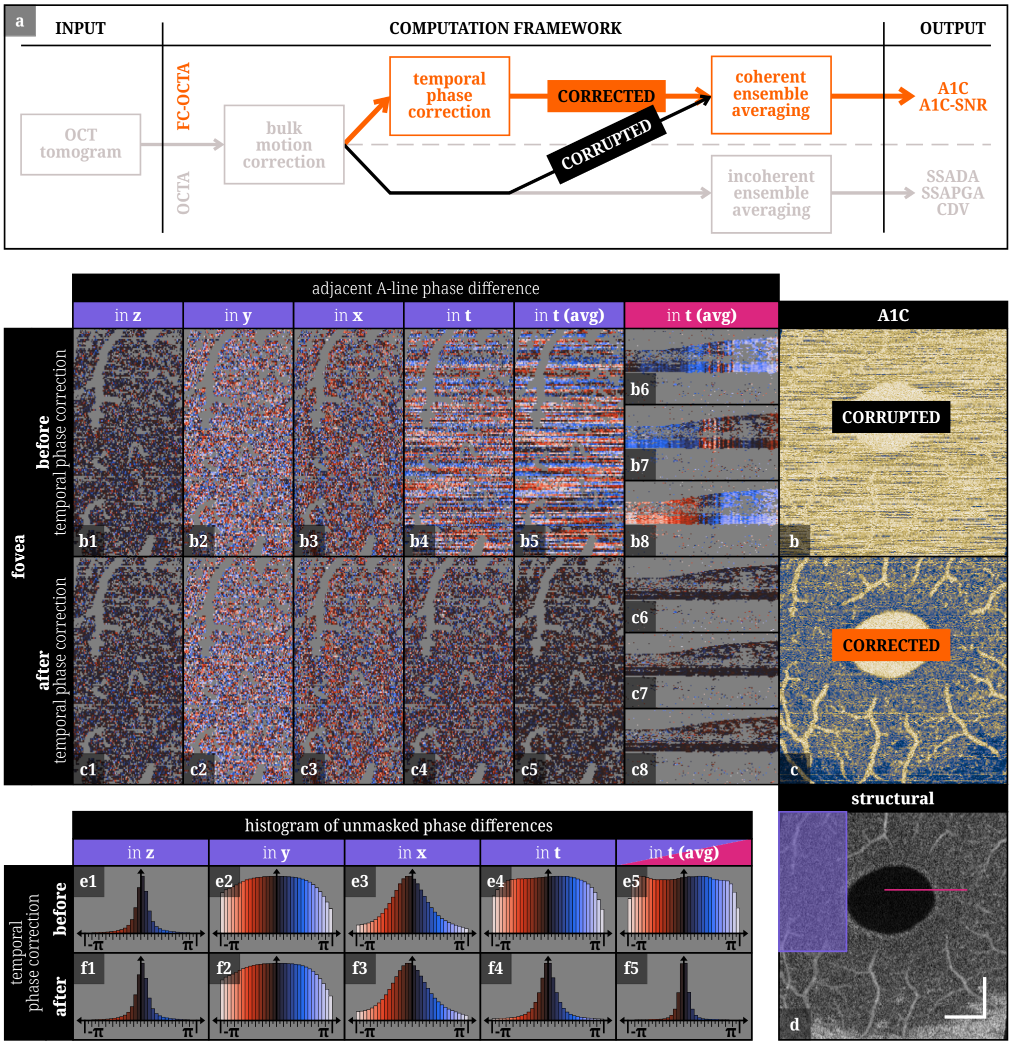


**Fig. S4.** (b1–b4, c1–c4) Adjacent A-line phase differences along all four spatiotemporal dimensions $(z, y, x, t)$ is presented in a cropped *en face* view [labeled as the purple rectangle in (d)] for the fovea of a healthy volunteer with and without temporal phase correction (voxels belonging to blood vessels and low-signal regions have been masked out in gray). (b5–b8, c5–c8) Additionally, the temporal phase difference for each repetition pair after 3D spatial ensemble averaging, $t$ (avg), is provided as a cropped B-scan cross-section [labeled as the pink line in (d)]. (b1, e1) Phase is intrinsically stable in $z$ in Fourier-domain OCT, which is demonstrated in the narrow, zero-mean phase difference distribution in the cropped *en face* view and histogram of unmasked phase differences. (e1–e5, f1–f5) These histograms were created using the entire volume and all temporal repetitions, ignoring only the grayed-out voxels. Each histogram corresponds to an exemplary *en face* view, with $t$ (avg) presenting (c6–c8) an additional B-scan cross-section to highlight the removal of phase noise in phase differences by performing temporal phase correction and the improvement in correction by incorporating 3D spatial ensemble averaging. The vertical axis of all histograms has been adjusted for visualization purposes. (b2–b3, c2–c3) Phase noise corrupts the phase difference between adjacent A-lines in both $x$ and $y$, which persists through temporal phase correction. However, after temporal phase correction, (f4, f5) the adjacent A-line phase difference for $t$ and especially $t$ (avg) approach (e1, f1) the distribution in $z$ for all spatiotemporal locations of static tissue, indicating the successful removal of temporal phase-noise differences, regardless of phase noise persisting along spatial dimensions. Thus, (a, b, c) temporal phase correction enables coherent 4D ensemble averaging and the calculation of fully-coherent OCTA metrics such as A1C. The white scale bars are 1 mm.

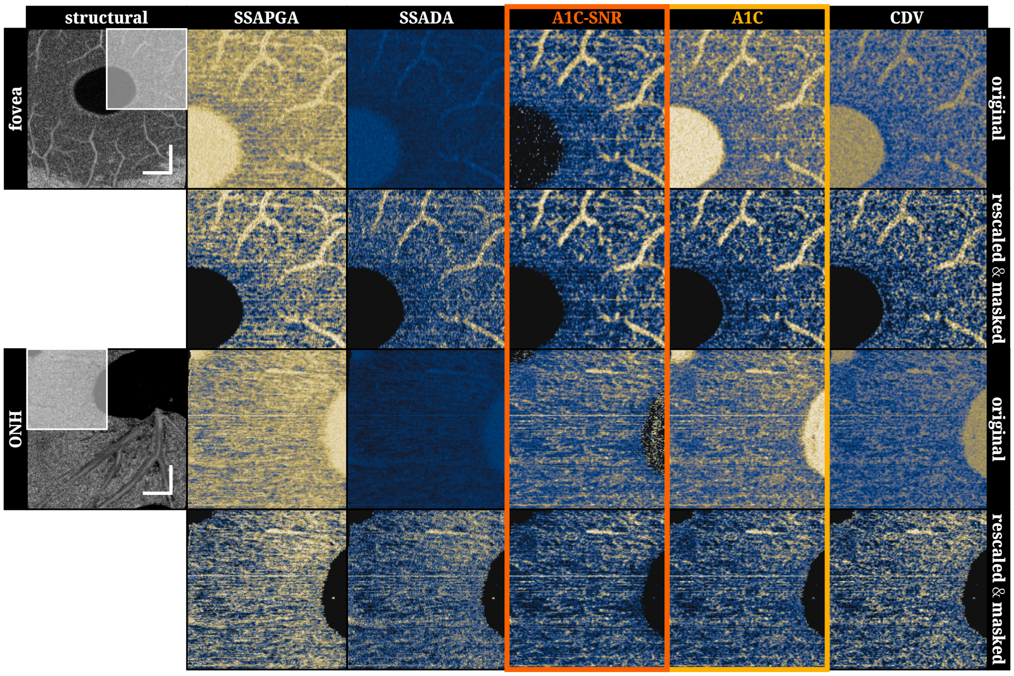


**Fig. S5.** Zoomed-in version of main article, Fig. 4. For both datasets, even after rescaling and SNR masking, A1C-SNR provides the best contrast between static tissue and blood vessels. By having greater dynamic range, more capillaries can be delineated in the fovea dataset, and the large vessel segment among the axonal bundles in the retinal nerve fiber layer is also more visible in the optic nerve head dataset. Vessel segments are observed to be shorter in the single-depth *en face* views of A1C-SNR and A1C compared to strong axial averaging methods SS-APGA and SSADA as less merging is performed along depth, making A1C-SNR and A1C better metrics for 3D microvasculature mapping and analysis.
